# Brain-behaviour relationships in the perceptual decision-making process through cognitive processing stages

**DOI:** 10.1101/2020.05.23.104620

**Authors:** Elaheh Imani, Ahad Harati, Hamidreza Pourreza, Morteza Moazami Goudarzi

## Abstract

Perceptual decision making, as a process of detecting and categorizing information, has been studied extensively over the last two decades. In this study, we investigated the neural characterization of the whole decision-making process by discovering the information processing stages. Such that, the timing and the neural signature of the processing stages were identified for individual trials. The association of stages duration with the stimulus coherency and spatial prioritization factors also revealed the importance of the evidence accumulation process on the speed of the whole decision-making process. We reported that the impact of the stimulus coherency and spatial prioritization on the neural representation of the decision-making process was consistent with the behavioral characterization as well. This study demonstrated that uncovering the cognitive processing stages provided more insights into the decision-making process.

## 1 Introduction

In daily work, people often encounter situations to select an action based on noisy sensory inputs. The process of choosing an action based on noisy sensory information is called perceptual decision making. Different cognitive processing stages are needed to receive the sensory information, accumulate perceptual evidence, and map to motor actions to accomplish the decision-making process in the brain [1, 2]. Various computational models have been proposed to describe the decision-making process at the behavioral level based on reaction time (RT) and response accuracy [3–6]. Accumulating the noisy perceptual evidence over time to reach the decision boundary prior to the response execution is the common idea across these models. Since this process is inherently noisy, the decision-making system needs some time to collect enough evidence to make a decision [7]. Recently, several researchers investigated the neural basis of the decision-making process. Initial analysis using single set recordings in monkeys during a random dot motion task discovered the neurons involved in this cognitive process [8–10]. These studies, inspired by the mathematical models of decision-making, demonstrated that neurons at the parietal and prefrontal cortex contributed to the decision-making process. These neurons integrate the information received from the sensory area, and the evidence accumulation is continued up to the fixed decision boundary.

Later approaches describe the decision-making process at both behavioral and physiological levels on human subjects with more complex tasks. The non-invasive imaging of human subjects, such as functional magnetic resonance imaging (fMRI) and magneto/electroencephalography (M/EEG), provided more extensive brain networks involved in the decision-making process compared to the single set recording of animal subjects. These approaches provided the possibility to characterize the decision-making process at a whole-brain level and clarified the neural correlate of decision making parameters with the physiological data [11, 12].

The studies in this area investigated the association between decision making parameters with fMRI and EEG data. The fMRI data with the high spatial resolution was employed to discover the brain regions that were functionally involved in the decision-making process [13–25]. Whereas, the EEG data with higher temporal resolution characterized this process with millisecond precision [21, 26–33]. The research on the fMRI data reported regions, such as the left dorsolateral prefrontal cortex [13], lateral occipital cortex [14], anterior Insula [15–17], inferior frontal sulcus [16], right Insula [18], right inferior frontal gyrus [19, 20], medial frontal gyrus [19], posterior-medial frontal cortex [21], and dorsomedial prefrontal cortex [20], responsible for evidence accumulation. Other studies reported the neural correlate of the decision boundary by changing the speed and accuracy tradeoff. They found that regions including the premotor area [22, 24, 25], striatum [24, 25], basal ganglia, thalamus, dorsolateral prefrontal [22], and dorsal anterior cingulate [23] had higher activations when preparing for speed rather than accurate decisions.

On the other hand, studies on the EEG data discovered the centro-parietal positivity (CPP) [26–29] and posterior parietal positivity [21] with gradually growing until response execution as the sensory accumulation process. Additionally, it was disclosed that the power of the parietal theta oscillations [30] and the beta oscillations of the motor cortex [31] covaried with the evidence accumulation. The studies of the neural basis of the decision boundary revealed that the power activity of the medial prefrontal cortex at theta frequency band associated with the value of decision boundary [32, 33].

While the decision-making process comprised of multiple processing stages, the previous research only described part of this process at the neural level. They sought to bridge the gap between brain and behavior by discovering the association between physiological data and components of the decision-making process, such as evidence accumulation. Characterizing the timing and neural signature of all the processing stages is necessary to provide a complete description of this process.

Recently some mathematical models were introduced to uncover the processing stages of the memory retrieval process [34, 35]. These approaches characterized the timing and neural signature of the processing stages of memory retrieval. However, in these studies, the memory retrieval process was described just at the physiological level. In our study, we take a similar approach to characterize the decision-making process at the physiological level and additionally clarify the association between the neural and behavioral characterizations. Using this approach, one can test the impact of a specific condition on the whole process of decision-making.

In this study, we utilized a recently published dataset of perceptual decision making [36], including both behavioral and physiological data. We employed a new approach to describe the decision-making process at both levels of brain and behavior and to uncover the relationships between them. Using this approach, we estimated the timing and brain signature of the processing stages. We assessed the effect of the internal subject state (spatial prioritization) and external world state (stimulus coherency) on the decision-making process at both behavioral and physiological levels. Finally, the relationships between brain and behavior were characterized in the decision-making process.

## 2 Materials and Methods

### 2.1 Experimental Design

In this study, a recently published decision-making dataset [36] was employed. This dataset includes both the physiological (EEG and fMRI) and behavioral (RT and response correctness) data from seventeen members of healthy adults aged 20-33 years. Participants categorized objects in a 2×2 factorial design task with the internal subject state (spatial prioritization) and external world state (stimulus coherency) factors (Figure 1). At each trial, the scrambled image of a car or a face was presented on the right or left visual hemifield for 200*ms*. Participants were asked to categorize objects as quickly and as accurately as possible by pressing their right or middle finger for selection in each category. For the first factor of the task design, the visual informativeness of the stimulus was manipulated by altering the phase of the images at the low and high coherency levels. For the prioritization factor, a cueing arrow indicating the visual hemifield of the stimulus was shown for 1000*ms* before the stimulus presentation on half of the trials. On the other half of the trials, a two-sided cueing arrow was presented for 1000*ms*. After the disappearing cueing arrow, the stimulus was presented randomly on each visual hemifield. These two factors created four different conditions: high coherency – prioritized (HCP), high coherency - not prioritized (HCNP), low coherency – prioritized (LCP), and low coherency – not prioritized (LCNP). For more information please refer to [36].

### 2.2 Data Acquisition

The data acquisition inside the magnetic resonance (MR) scanner with simultaneous recording of EEG and fMRI included 90 trials for each condition with an intertrial interval of 10 to 12 seconds. These 90 trials were split into five separate experimental sessions. All EEG data were acquired with a 64-channel MR compatible EEG system. The scalp electrodes on the EEG cap follow the 10-20 system in naming and placement, which include two additional channels, one for recording the electrocardiogram (ECG) and the other for recording the electrooculography (EOG).

### 2.3 Data Preprocessing

We employed the re-referenced and MR-related artifact-free version of EEG data, which was provided by the owners. The EEG data was downsampled to 500Hz. Further data preprocessing was performed in this study using the Fieldtrip toolbox [37]. EEG data was bandpass filtered (a zero-phase, two-pass, and fourth-order Butterworth filter) from 1HZ to 70HZ followed by band-stop filtering (a zero-phase, two-pass, and fourth-order Butterworth filter) from 48HZ to 52HZ. The significant artifactual EEG sections were selected visually and were ignored before further analysis. The eye movement and muscle artifacts were removed using an independent component analysis (ICA) algorithm. The artifactual components were selected visually and rejected from the components set, and the artifact-free data was obtained by reconstructing the refined ICA components. The stimulus-lock epochs with the length of RT were extracted from EEG data and were baseline corrected using the 200*ms* pre-cue baseline (the time interval between 1200*ms* to 1000*ms* pre-stimulus). Of the initial total of 16 participants (one subject did not have behavioral data) with five sessions for each subject, four subjects were removed from further analysis because of the significantly poor EEG data quality. From the remaining subset, a total of 13 sessions were rejected as well.

**Figure 1:**
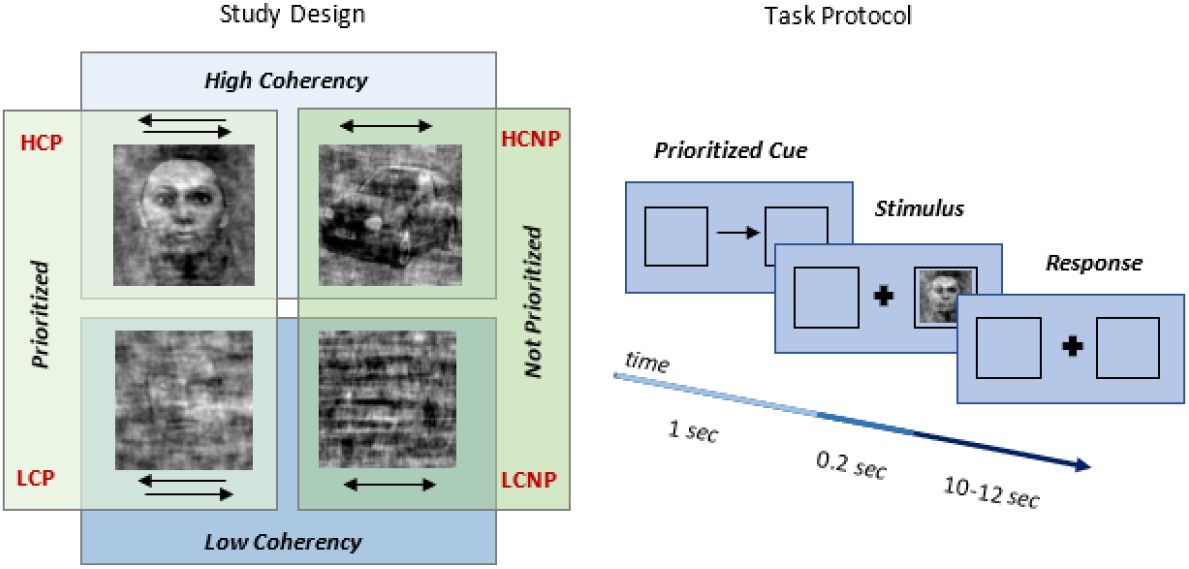
Task protocol overview. A 2×2 factorial design task with spatial prioritization and stimulus coherency factors. At each trial, the scrambled image of a car or a face was presented on the right or left visual hemifield for 200*ms*. For the coherency factor, the visual informativeness of the stimulus was manipulated by altering the phase of the images at low and high coherency levels. For the prioritization factor, a cueing arrow indicating the visual hemifield of the stimulus was shown for 1000*ms* before the stimulus presentation on half of the trials.

### 2.4 Behavioral Data Analysis

The drift-diffusion model (DDM) as a most discussed model for evidence accumulation process, was employed to analyze the decision-making process at the behavioral level. The DDM views decision making as a process of noisy accumulation of evidence over time (Figure 2), which is parameterized by a set of three parameters, i.e., drift rate, decision boundary, and bias. The average rate of accumulating the noisy evidence was called drift rate, *v*, which models the efficiency of the evidence accumulation. Such that, more efficient evidence accumulation leads to higher drift rates [38]. The higher drift rate is also associated with a faster decision-making process [7]. In this model, the sensory evidence from perception accumulated over time until it reached a decision boundary, *a,* for each choice. The decision boundary shows the amount of evidence is needed to be accumulated after stimulus encoding until response onset. Models sometimes include a bias parameter, *z*, when there is some prior knowledge about the task. This model can separate the decision component from non-decision ones, such as stimulus encoding and response execution [7, 39]. These non-decision components together have a mean-time, *t*_*er*_, which is called non-decision time. The decision component is also characterized by the drift rate parameter.

In this study, we evaluated the decision-making performance under different conditions. The hierarchical DDM (HDDM) [40] was employed to estimate model parameters *v*, *a* and *t*_*er*_. Markov chain Monte Carlo (MCMC) sampling was employed to approximate the posterior probability of the model parameters at the individual and group levels. We initialized the HDDM with 10000 posterior samples and discarded the first 1000 samples as burn-in. To show the effect of the coherency and spatial prioritization on the decision-making performance, the parameters of the HDDM were estimated separately for different conditions (HCP, HCNP, LCP, and LCNP). A two-way repeated measure ANOVA was then employed to test the effect of stimulus coherency and spatial prioritization on the decision and non-decision components of the DDM.

**Figure 2:**
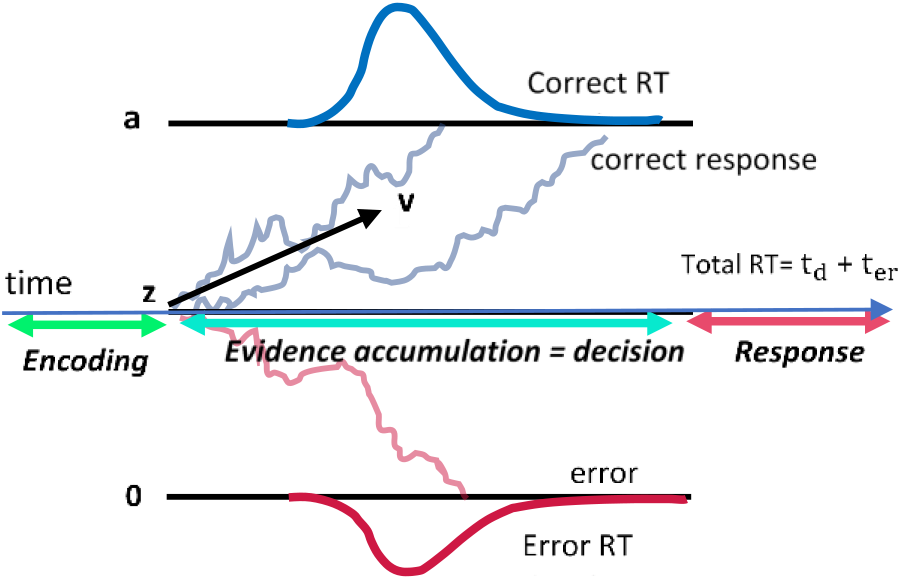
The diffusion decision model. Noisy evidence is accumulated from the starting point, *z,* over time (during *t*_*d*_ *ms*) with the average drift rate, *v,* until it reaches the decision boundary, *a*. The non-decision components such as encoding and response output have a mean time called non-decision time, *t*_*er*_. Thus, total RT includes non-decision time, *t*_*er*_, and decision time, *t*_*d*_.

### 2.5 EEG Data Analysis

As mentioned before, the decision-making process consists of some processing stages, i.e., encoding, decision making, and response execution. At the behavioral level, the encoding and response execution together were considered as a single non-decision component, which was characterized by the non-decision time parameter. The decision stage was also viewed as an evidence accumulation process that was described by the drift rate parameter. Uncovering the timing and the neural basis of these processing stages provides more insight into this process. In this study, we characterized the decision-making building blocks by employing the HSMM-EEG method [34] at the physiological level. This method is based on a hidden semi-Markov model that assigns each sample to one stage and determines the timing of the transition between the stages (Figure 3). Using this method, each stage is characterized by duration and signature parameters. To reduce the model complexity and to preserve the temporal profile of the EEG data to the HSMM, instead of EEG samples, the windowed EEG signal naming snapshot was employed in this model. For more details please see [34]. Finally, the HSMM-EEG was fitted using the denoised low dimensional representation of the snapshots. The dimensionality reduction was performed by a principal component analysis (PCA), which accommodated 98% of the variance of the data.

**Figure 3:**
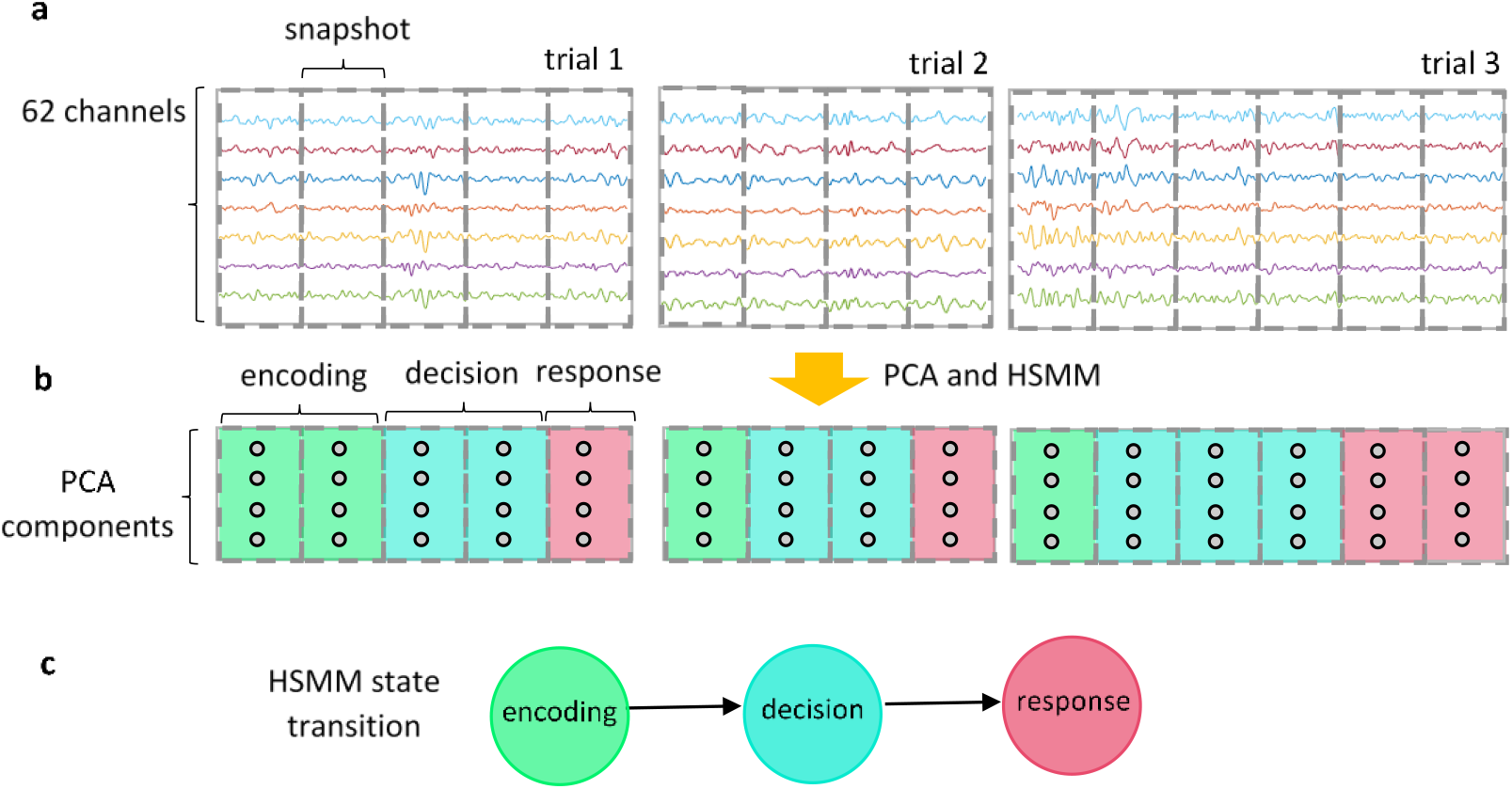
Information processing stages extraction using HSMM-EEG analysis. **a.** Shows windowing EEG signals to create snapshots. **b.** The snapshots are preprocessed to reduce dimensionality. The preprocessed snapshots are fed to the HSMM-EEG to separate decision-making stages. The results of the HSMM-EEG on the snapshots are shown with three colors for encoding (green), decision (blue), and response (red). **c.** Illustration of the HSMM-EEG stage transition with encoding, decision, and response stages.

Using a snapshot length of 160*ms* with 101 PCA components, which accounted for 98% of the total data variance, the data was analyzed to uncover three stages of the decision-making process. The two-way repeated measure ANOVA was then employed to test the association between coherency and prioritization factors and the duration of the stages. Relationships between behavioral and neural levels were also investigated using Pearson’s correlation analysis.

## 3 Results

### 3.1 Decision-making characterization at the behavioral level

The coherency of the stimulus was changed at two different levels to influence the amount of sensory evidence. The spatial prioritization factor was also applied to change the internal subject state. The participants were asked to categorize cars versus faces under the combination of these factors. The response time and response accuracy were analyzed to check the influence of these two factors on decision-making performance. The results of the two-way repeated measure ANOVA revealed the significant main effect of the coherency factor on the response time (*F*_(1,31)_ = 41.61, *p* < 0.001) and response accuracy (*F*_(1,31)_ = 130.23, *p* < 0.001). Such that, increasing the coherency level of the stimulus provided faster and more accurate responses (Figure 4). The spatial prioritization had also a significant main effect on the response time (*F*_(1,31)_ = 5.48, *p* = 0.025), and marginally significant effect on the response accuracy (*F*_(1,31)_ = 3.92, *p* = 0.056).

**Figure 4:**
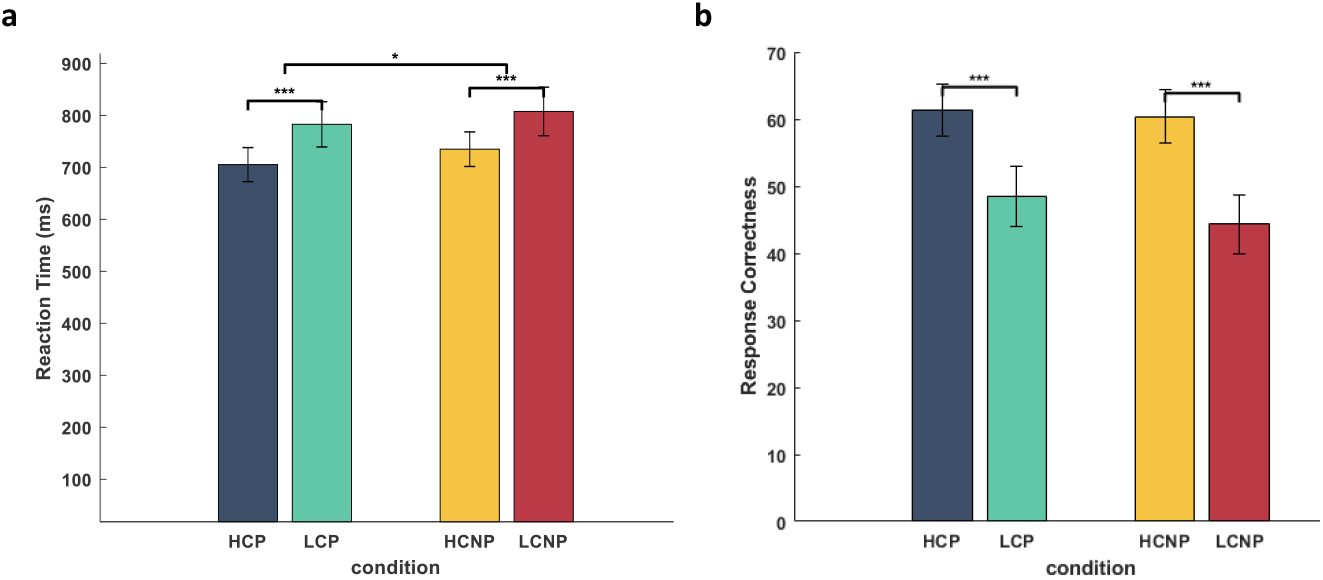
mean response time and response accuracy across subjects. **a.** demonstrates the mean response time across subjects for each condition. The two-way repeated measure ANOVA revealed the significant main effect of the coherency (*F*_(1,31)_ = 41.61, p < 0.001) and spatial prioritization factors (*F*_(1,31)_ = 5.48, p = 0.025) on the response time. b. illustrates the mean response correctness across subjects for each condition. The results show a significant main effect of coherency (*F*_(1,31)_ = 130.23, p < 0.001) and a marginally significant main effect of spatial prioritization (*F*_(1,31)_ = 3.92, p = 0.056) on the response accuracy. The sign ‘***’ and ‘*’ symbolize p-value < 0.001 and p-value < 0.05.

To further characterize the influence of the coherency and spatial prioritization on the decision-making process, the decision and non-decision components of this process were estimated using DDM. We hypothesized that the decision component of the process was the only processing stage associated with stimulus coherency and spatial prioritization factors. We also showed that the coherency factor had a stronger relationship with evidence accumulation than the prioritization factor.

To test this hypothesis at the behavioral level, the drift rate and non-decision time parameters of the DDM were estimated for each condition (HCP, HCNP, LCP, and LCNP). These two parameters characterized the decision and non-decision components of the process, respectively. The results of the statistical test demonstrated that coherency significantly affected the drift rate parameter, as determined by two-way repeated measure ANOVA (*F*_(1,31)_ = 294.65, *p* < 0.001) (Figure 5). Moreover, we found a marginally significant effect of the prioritization factor on the drift rate (*F*_(1,31)_ = 4.09, *p* = 0.052). As expected, the results revealed the major impact of the coherency factor on the efficiency of evidence accumulation rather than the spatial prioritization factor. Additionally, we checked whether the coherency and spatial prioritization were associated with the non-decision component. As depicted in Figure 5, the two-way repeated measure ANOVA disclosed no significant main effect of the stimulus coherency (*F*_(1,31)_ = 0.62, *p* = 0.44) and spatial prioritization (*F*_(1,31_) = 0.08, *p* = 0.78) factors on the non-decision time parameter.

Overall, the results demonstrated that among the different processing stages of decision making, only the evidence accumulation covaried with stimulus coherency. The spatial prioritization had a marginally significant impact on the decision stage as well. Since the non-decision component was not associated with factors of interest, the efficiency of the decision-making process was more dependent on the evidence accumulation stage.

**Figure 5:**
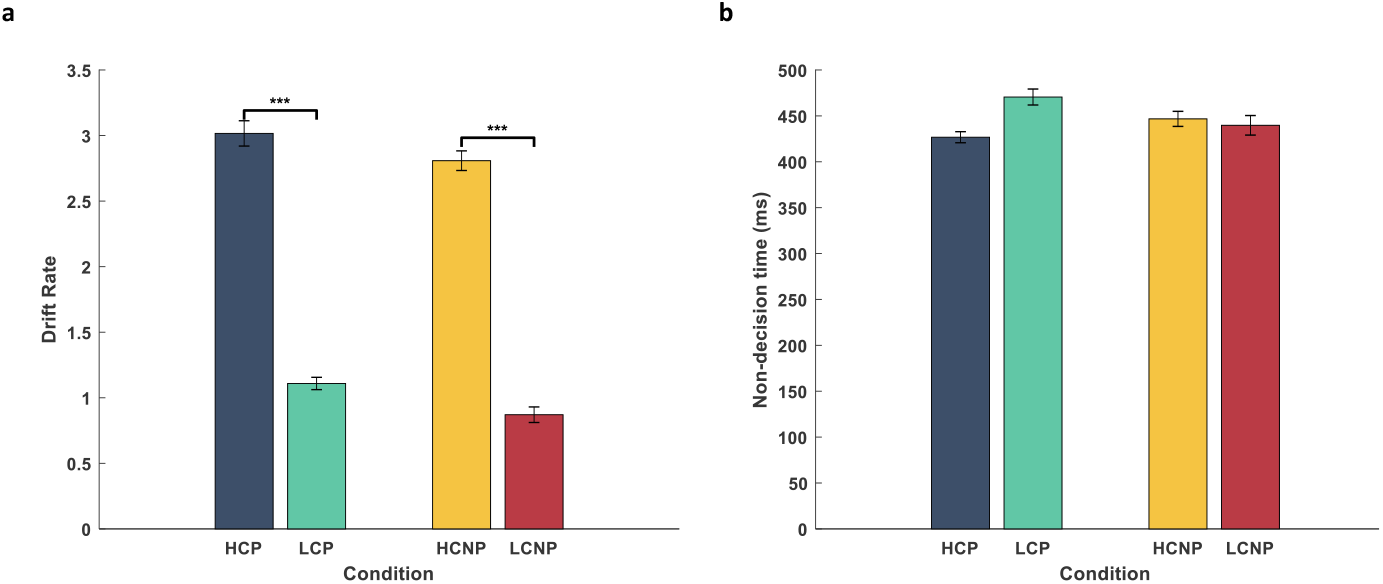
The predicted drift rate and non-decision time. **a.** Illustration of the mean of the drift rate parameter across subjects for different conditions (HCP, HCNP, LCP, and LCNP). The significant main effect of the coherency factor was reported by two-way repeated measure ANOVA (*F*_(1,31)_ = 294.65, *p* < 0.001). Moreover, a marginally significant difference was found by the prioritization factor by two-way repeated measure ANOVA (*F*_(1,31)_ = 4.09, *p* = 0.052). **b.** Presentation of the mean of the non-decision time parameter across subjects for each condition. There was no significant main effect of coherency (two-way repeated measure ANOVA, *F*_(1,31)_ = 0.62, *p* = 0.44) and prioritization factors (two-way repeated measure ANOVA, *F*_(1,31)_ = 4.09, *p* = 0.052) on this parameter. Error bars represent standard error of the mean. The sign ‘***’ symbolizes *p*-value < 0.001.

### 3.2 Decision-making characterization at the neural level

Next, we investigated whether we could characterize the decision-making process in more detail by estimating the timing and neural signature of the individual processing stage. With the benefit of EEG data with millisecond temporal resolution and the use of the HSMM-EEG method, we extracted the timing of each processing stage for each condition, i.e., HCP, HCNP, LCP, and LCNP (Figure 6). Since the sequences of the stages were the same for all conditions, the conditions were analyzed jointly with HSMM-EEG. Using the duration of the processing stages, we tested the previous hypothesis and checked whether coherency and spatial prioritization were only associated with decision stage duration. As shown in Figure 6, the two-way repeated measure ANOVA found no significant main effect of stimulus coherency (*F*_(1,31)_ = 0.61, *p* = 0.44) and spatial prioritization (*F*_(1,31)_ = 0.9, *p* = 0.36) factors on the encoding stage duration. As expected, the coherency had a significant main effect on the duration of the decision stage as determined by two-way repeated measure ANOVA (*F*_(1,31)_ = 24.28, *p* < 0.001). However, the two-way repeated measure ANOVA disclosed no significant effect of spatial prioritization (*F*_(1,31)_ = 2.48, *p* = 0.13) on the duration of this stage. The response execution stage was also analyzed by the two-way repeated measure ANOVA to find the impacts of coherency and prioritization on the duration of this stage. The results of the statistical test revealed no significant association between coherency (*F*_(1,31)_ = 3.96, *p* =0.06) and spatial prioritization (*F*_(1,31)_ = 0, *p* = 0.95) factors on response duration.

The investigation of the decision-making process at the physiological level also confirmed the findings of behavioral analysis. The results demonstrated that decision making is affected by the coherency factor rather than spatial prioritization. Changing the coherency level of the stimulus had only association with the decision stage duration. It might be because stimulus coherency affected the input sensory evidence quality. Therefore, lower stimulus coherency needed more time to reach the decision boundary, which was characterized by the evidence accumulation process.

**Figure 6:**
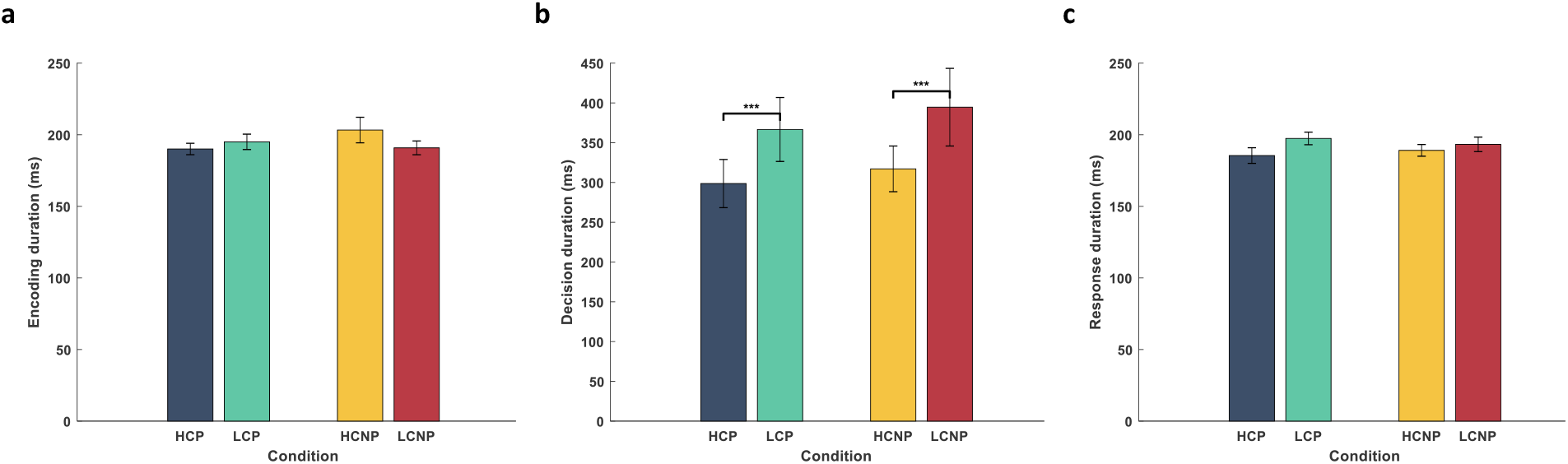
Estimated processing stages duration. Illustration of the mean timing of decision-making stages, i.e., encoding, decision, and response for individual condition (HCP, HCNP, LCNP, and LCNP). The two-way repeated measure ANOVA found no significant main effect of coherency (*F*_(1,31)_ = 0.61, *p* = 0.44) and spatial prioritization (*F*_(1,31)_ = 0.9, *p* = 0.36) on the duration of the encoding stage. The significant effect of the coherency was found by the two-way repeated measure ANOVA on the decision stage duration (*F*_(1,31)_ = 24.28, *p* < 0.001). However, there was no significant association between the decision stage and the prioritization factor (*F*_(1,31)_ = 2.48, *p* = 0.13), as determined by ANOVA. Finally, no significant effect of coherency (*F*_(1,31)_ = 3.96, *p* =0.06) and prioritization (*F*_(1,31)_ = 0, *p* = 0.95) was found by the two-way repeated measure ANOVA on the response duration. Error bars represent standard error of the mean. The sign ‘***’ symbolizes *p*-value < 0.001.

To further characterize the decision-making process, the signature of the states were computed by taking the weighted average of the snapshots across trials. The weight of each snapshot was the probability of belonging that snapshot to each state which was estimated by the HSMM-EEG. The resulted signatures are illustrated in Figure 7. As shown in this figure, the occipital negativity was observed at the encoding stage. It was consistent with previous findings that the posterior visual N200 component of the event-related potential (ERP) was associated with the sensory processing [41–43]. At the next two stages, the encoded evidence accumulated over time to reach the decision boundary and finalize the response execution. The signature of the decision and response stages revealed the increase at the CPP at the decision and response execution stages. Previous research also disclosed the increasing of the CPP with incoming evidence that peaked at the time of response execution [26, 29]. The neural basis of response execution clarifies the positivity of the left motor cortex as well. It might be because as most of the subjects are right-handed (9 of 11 subjects), the activity of the left motor cortex was increased at the response execution stage.

**Figure 7:**
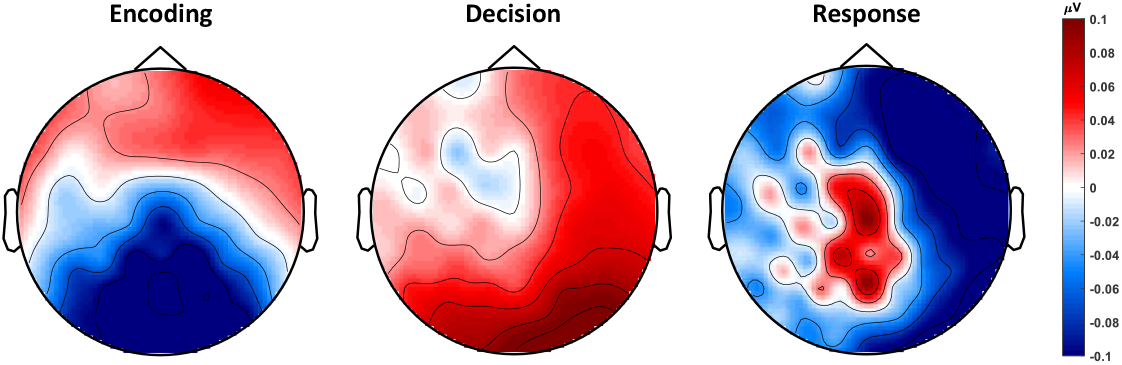
Signature of the processing stages. The signatures were created by taking the weighted average of the snapshots across trials. **a.** The signature of the encoding stage disclosed the negativity of the visual areas. **b** and **c** illustrate the decision and response stages, respectively. The signature of these two stages demonstrated the increase of CPP until response execution. The positivity of the left motor cortex was also depicted at the response execution stage.

### 3.3 Brain-behavior relationships

We investigated the association between behavioral and neural representations of the decision-making process. At the behavioral level, the DDM considers the drift rate and non-decision time parameters as a measure to describe the decision and non-decision stages of the process. Additionally, the HSSM-EEG method decomposes the decision-making process into its components at the physiological level by estimating the timing of each processing stage. To probe the relationships between decision-making characterization at the physiological and behavioral levels, we examined the association between DDM parameters and the duration of the information processing stages.

As reported with previous research, the faster decisions associated with a higher drift rate [7]. Thus, we investigated whether the drift rate parameter had a negative interaction with the duration of the decision stage at the physiological level. Figure 9a indicates the estimated drift rate and duration of the decision stage for each condition. A significant negative correlation (*r* = −0.47, p = 0.001, Pearson’s correlation) was found between the physiological and behavioral characterizations of the decision stage, which was consistent with the previous findings [7]. Additionally, we checked whether the non-decision parts of the decision process had any interaction between brain and behavior representations. Figure 9b shows the non-decision time parameter estimated by the DDM and the sum of the duration of encoding and response stages resulted from the HSSM-EEG model for each condition. The results demonstrated a significant positive interaction (*r* = 0.78, *p* < 0.001, Pearson’s correlation) between behavioral and physiological representations of the non-decision component.

We further examined the relationships between physiological representation of the decision-making process and the RT. We hypothesized that the duration of the whole decision-making process depended more on the decision stage rather than the non-decision stages. To test this hypothesis, the interaction between each stage duration with the RT was estimated using Pearson’s correlation across subjects and conditions (Figure 9). A significant interaction was found between the duration of the decision stage and RT (*r* = 0.99, *p* < 0.001, Pearson’s correlation). The correlation analysis revealed no significant interactions between RT and duration of the encoding (*r* = 0.25, *p* = 0.11, Pearson’s correlation) and response (*r* = 0.29, *p* = 0.06, Pearson’s correlation) stages. The results disclosed that the most important factor in RT is the duration of the evidence accumulation process. The faster accumulation process led to shorter reaction times.

**Figure 8:**
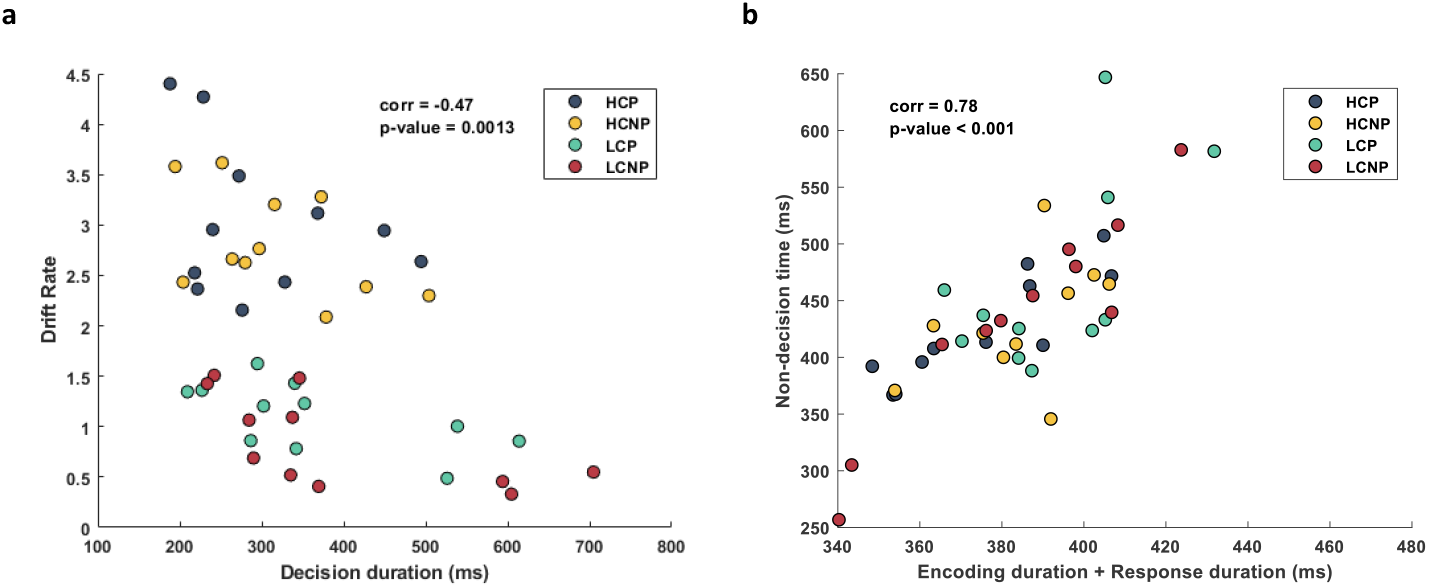
Association between behavioral and physiological representations of the decision-making process. **a.** Shows the interaction between the drift rate parameter of the DDM and the decision stage duration of the HSMM-EEG model for each condition. The significant negative interaction (*r* = −0.47, p = 0.001, Pearson’s correlation) was found between these parameters. **b.** Illustrates the non-decision time and sum of encoding and response stages duration for each condition. The results clarified a significant positive correlation (*r* = 0.78, *p* < 0.001, Pearson’s correlation) between these parameters.

**Figure 9:**
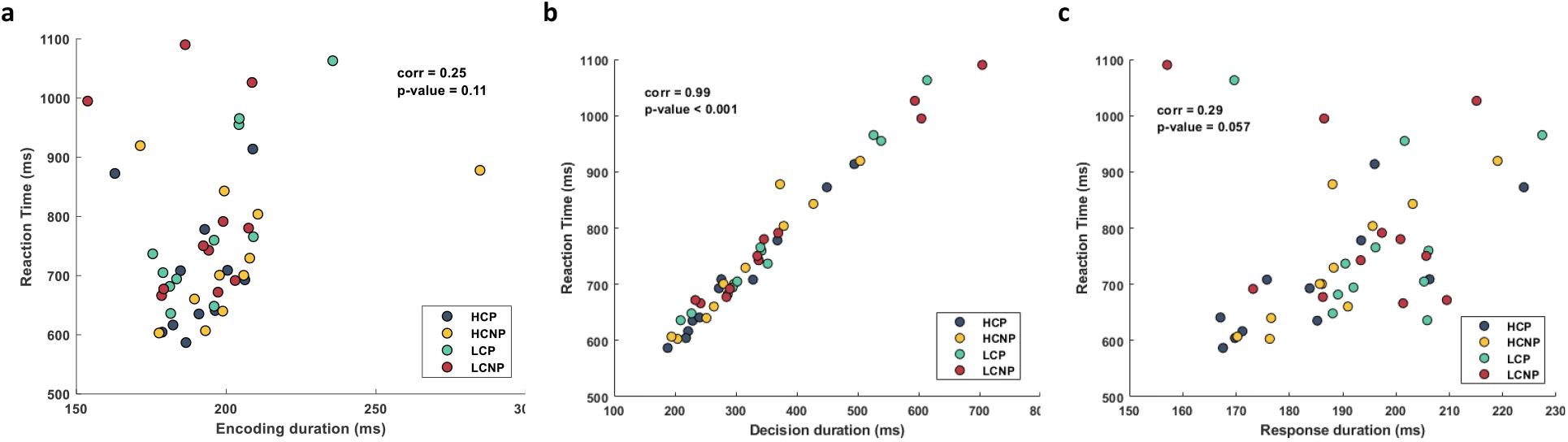
Relationships between reaction time and decision stages duration. Panels a, b, and c depict the mean value of the RT and the encoding, decision, and response duration respectively for each subject and condition. A significant interaction was found between the RT and duration of encoding (*r* = 0.25, *p* = 0.11, Pearson’s correlation), decision (*r* = 0.99, *p* < 0.001, Pearson’s correlation), and response (*r* = 0.29, *p* = 0.06, Pearson’s correlation) states.

## 4 Discussion

The classical model of the decision-making process encompasses three distinct processing stages, i.e., encoding, decision, and response execution. Here we shed light on the decision-making process by characterizing/linking this process at both neural and behavioral levels. At the neural level with the aim of EEG data with a high temporal resolution, the timing and neural signature of the processing stages of the decision-making process (encoding, decision, and response execution) were estimated. Taken together, the relationship between the behavioral data and different stages of the decision-making process on EEG segments provides insights into the underlying process.

Using the HSMM-EEG model, the stages of the decision-making process were extracted with each associated with the duration and neural signature parameters (Figure 6 and Figure 7). The duration of the encoding stage depicted in Figure 6a revealed that the time needed for sensory inputs to be encoded was nearly 190*ms*. This is consistent with the previous study, which clarified that the latency of the N200 component of the ERP reflected the time of sensory encoding [43]. The previous animal studies also revealed that the lateral inter-parietal (LIP) neurons started to accumulate the perceptual evidence about 200*ms* after stimulus presentation [8]. During the encoding stage, the occipital negativity was observed in the early visual area as well (Figure 7a). As reported by recent studies [41–43], in line with our result, the N200 component at the visual area associated with perceptual processing. The MEG study on the frequency domain also disclosed the gamma band activity on the visual cortex as the stimulus encoding [2].

Next, the encoded sensory inputs were accumulated over time to reach the decision boundary at the decision stage. The neural signature of the decision stage revealed the increase of the CPP decision and response stages (Figure 7b,c). Similarly, previous studies found that CPP was associated with the evidence accumulation process with a pick at the time of response execution [26–29], which confirmed the resultant signatures.

When the accumulated perceptual evidence reached the decision boundary, the motor execution initiated. As shown in the signature of the response stage (Figure 7c), besides increasing the CPP, the positivity of the left motor cortex was observed as well. It was because most of the participants were right-handed (9 of 11 subjects). The processing at the motor execution stage, on average, lasted about 190*ms,* as depicted in Figure 6c. The study on the MEG data was also revealed that event-related desynchronization (ERD) peaked about 170*ms* before response execution over the sensorimotor area contralateral to the response side [44].

Furthermore, we hypothesized that the efficiency of the decision-making process was more dependent on the decision stage rather than on non-decision ones. Since the characterization of the decision-making process at the neural level provided the timing of the processing stages, we tested this hypothesis at both neural and behavioral levels. The statistical analysis on the physiological level disclosed that changing the coherency level of the stimulus associated with the decision stage duration. Such that, the higher coherent stimulus led to a shorter decision stage. Furthermore, the behavioral analysis provided similar results and revealed the significant interaction of stimulus coherency with the drift rate parameter, that confirmed the role of the drift rate as a perceptual input quality measure, as explained by the DDM [45]. The analysis of the single set recording also supported this finding and showed that faster responses were associated with the rapid build-up of LIP activity [8]. We also analyzed the relationships between the duration of processing stages and RT (Figure 9). The significant association between RT and the duration of the decision stage demonstrated the importance of this stage on the speed of the decision-making process.

According to the similar operation of the neural and behavioral representations of the decision-making process under task conditions, we test whether there was a relationship between these two levels (Figure 8). Consequently, we hypothesized that the drift rate parameter that characterizes the efficiency of the evidence accumulation process has an interaction with the duration of the decision stage at the physiological level. Such that, higher drift rates provide shorter decision stages. Similarly, we test whether the duration of the combination of the encoding and response stages have any interaction with the non-decision time of the DDM. To test these hypotheses, Pearson’s correlation was conducted, and the results confirmed the significant association between physiological and behavioral levels. The DDM also considered the drift rate as a speed of evidence accumulation. Such that faster decisions related to the higher drift rates [7]. These findings verified the results of this study, as well.

## 5 Conclusion

In this study, we sought to bridge the gap between neural and behavioral representations of the decision-making process. Neural characterization of the decision-making process was uncovered through information processing stages. We showed that these two representations had a similar manner under different stimulus coherency levels. Additionally, the results at both neural and behavioral levels revealed the importance of the decision stage on the efficiency of the whole decision-making process. The significant association between processing components at both neural and behavioral levels was also a validation for the neural characterization of the decision-making process. Overall results demonstrated that this representation provides more insight into the decision-making process by providing both the duration and neural signature of the cognitive stages.

## Acknowledgments

All data used in this research is publicly available in the open science framework (https://osf.io/). We would like to thanks Dr. Dirk Ostwald and his colleagues that make this data publicly available.

